# Antigen-induced but not innate memory CD8 T cells express NKG2D and are recruited to the lung parenchyma upon viral infection

**DOI:** 10.1101/224782

**Authors:** Morgan Grau, Séverine Valsesia, Julien Mafille, Sophia Djebali, Martine Tomkowiak, Anne-Laure Mathieu, Daphné Laubreton, Simon de Bernard, Pierre-Emmanuel Jouve, Laurent Buffat, Thierry Walzer, Yann Leverrier, Jacqueline Marvel

**Author notes:** Corresponding author: Jacqueline Marvel, Team Immunity and cytotoxic lymphocytes, CIRI, Inserm, U1111, Université Claude Bernard Lyon 1, CNRS, UMR5308, École Normale Supérieure de Lyon, Univ Lyon. 21 Avenue Tony Garnier - 69365 LYON cedex 07, France.

## Abstract

The pool of memory-phenotype CD8 T cells is composed of antigen-induced (AI) and cytokine-induced innate (IN) cells. IN have been described as having similar properties to AI memory cells. However, we found that pathogen-induced AI memory cells can be distinguished from naturally-generated IN memory cells by surface expression of NKG2D. Using this marker, we described the increased functionalities of AI and IN memory CD8 T cells compared to naive cells, as shown by comprehensive analysis of cytokine secretion and gene expression. However, AI differed from IN memory CD8 T cells by their capacity to migrate to the lung parenchyma upon inflammation or infection, a process dependent on their expression of ITGA1/CD49a and ITGA4/CD49d integrins.

## Introduction

One hallmark of the adaptive immune system is its ability to respond faster and stronger to previously encountered antigens (Ag). This immunological memory relies on the generation of cells that display increased reactivity towards the previously encountered Ag. Protection against intracellular pathogens or tumor-derived Ag is conferred in part by Ag-induced memory CD8 T cells (AI). Indeed, AI memory CD8 T cells have improved functional properties compared to naive cells, making them more potent to rapidly eliminate infected cells upon re-infection ^1,2^. These AI memory cells are found in secondary lymphoid organs, but a subset of them, the tissue-resident memory cells (TRM), settles within non-lymphoid tissues where they provide increased protection against secondary pathogen infections ^3,4^ TRM are long-lived sessile cells in most tissues except the lung where they need to be replenished from the circulating pool of memory cells ^5,6^

Memory-phenotype CD8 T cells or innate memory cells (IN) can also be generated through several alternative pathways that are independent of foreign Ag exposure ^7^. Memory CD8 T cells generated through lymphopenia-induced proliferation (LIP) were the first IN memory cells to be described ^8–12^. This pathway depends on strong IL-7 stimulation of naive CD8 T cells (due to the increased availability of this γc cytokine in the lymphopenic host) combined to weak TCR stimulation through self-peptide/MHC-complexes ^13–15^. Other γc cytokines also support the generation of IN memory cells. *In vivo*, strong IL-2 stimulation through injection of IL-2/anti IL-2 antibody complexes, was shown to drive the generation of IN memory CD8 T cells from naive TCR transgenic cells ^16^. Similarly, the characterization of several mutant mouse strains revealed that strong IL-4 stimulation of CD8 single positive thymocytes or naive CD8 T cells leads to IN memory cell generation ^17–19^. Moreover, naive BALB/c mice have an increased proportion of memory phenotype CD8 T cells due to higher levels of circulating IL-4 compared to naive C57BL/6 mice ^20^. Conversely, naive mice deficient for IL-4 production or signaling have a reduced frequency of IN memory CD8 T cells ^21,22^. In physiological conditions, LIP memory cells generation occurs during the neonatal period in naive mice ^21,23,24^ and Th2 immune responses might also favor the generation of IN memory CD8 T cells ^25^.

Hence, among CD8 T cells specific for foreign Ag never encountered by Specific Pathogen Free (SPF) naive mice, 10-20% of cells display a memory phenotype. These cells are also refered to as virtual memory cells ^26^. Importantly, equal numbers of these virtual memory cells were found in naive germ free mice, indicating that their generation is independent of microbiota-derived Ag ^26^. Therefore, in physiological conditions, the pool of memory-phenotype CD8 T cells is composed of two classes of cells: AI and IN.

AI and IN CD8 memory cells are generated through distinct pathways, but express a similar array of surface markers, which has hampered their demarcation. Interestingly, experimentally generated TCR transgenic OTI memory CD8 T cells do not express CD49d compared to AI OTI memory CD8 T cells. This lack of CD49d expression has been used to identify and characterize OVA and VV-specific clones of IN memory cells generated in physiological conditions ^26^. In parallel, it has been shown that compared to naive cells, these unconventional Ag-specific memory cells (IN or virtual memory cells) are able to mount an efficient response against pathogen infection with increased functional properties including augmented IFN-γ production and proliferative response ^18,26,27^. However, the comparison between IN and AI memory CD8 T cells in terms of phenotype, function and gene-expression profile has not been performed.

Upon strong TCR triggering, CD8 T cells express high levels of the NK cell receptor NKG2D ^28^ and antigen-induced memory cells express NKG2 ^29^. In contrast, IL-4-induced innate cells do not express NKG2D ^29^. Therefore, we hypothesized that NKG2D could be differently expressed between AI and IN memory CD8 T cell populations.

We herein demonstrate that the expression of NKG2D is restricted to AI memory CD8 T cell populations. Using NKG2D as a marker of AI cells, we performed an extensive comparison of AI and IN cells within the natural pool of memory CD8 T cells. Our results indicate that although IN CD8 T cells share many features with AI memory cells, only AI cells are recruited towards the lung parenchyma upon inflammation or infection.

## Results

### NKG2D expression identifies polyclonal antigen-induced memory CD8 T cell populations

We tested the hypothesis that NKG2D expression could discriminate Antigen-Induced (AI) from Innate (IN) memory cells. Indeed, IN cells generated following injection of IL-2 or IL-4 antibody complexes (Figure 1A and S1) or after lymphopenic proliferation (Figure S2) do not express NKG2D at their cell surface. In agreement with this hypothesis, in non-immunized SPF mice, the majority (more than 90%) of splenic memory-phenotype CD8 T cells (i.e. CD44^hi^) which are mainly IN cells ^7^ do not express NKG2D (Figure 1B). Similarly, virtual memory cells are mainly NKG2D negative (Figure 1C). In contrast, almost all B8R-specific memory CD8 T cells (CD44^hi^ B8R+) from C57BL/6 mice infected with vaccinia virus (VV) express NKG2D (Figure 1D). Following infection with VV, the frequency of CD44^hi^ effector or memory CD8 T cells expressing NKG2D was increased by more than a hundred-fold in the effector phase leading to a ten fold expansion of the NKG2D+ subset in the memory phase (Figure 1E and S3). This was not specific to VV as 55 days after infection by Influenza virus (Flu) or bacteria *Listeria monocytogenes* (Lm) there was also a strong amplification of the NKG2D positive CD44^hi^ CD8 T population in the blood (Figure 1E). In contrast, the number of memory-phenotype NKG2D− cells remained stable when comparing the naive and the memory phase. These results could be extended to other mouse strains: in BALB/c mice and in the outbred mouse strain OF1 VV infection induced the generation of NKG2D+ memory CD8 T cells (Figure S4).

**Figure 1:**
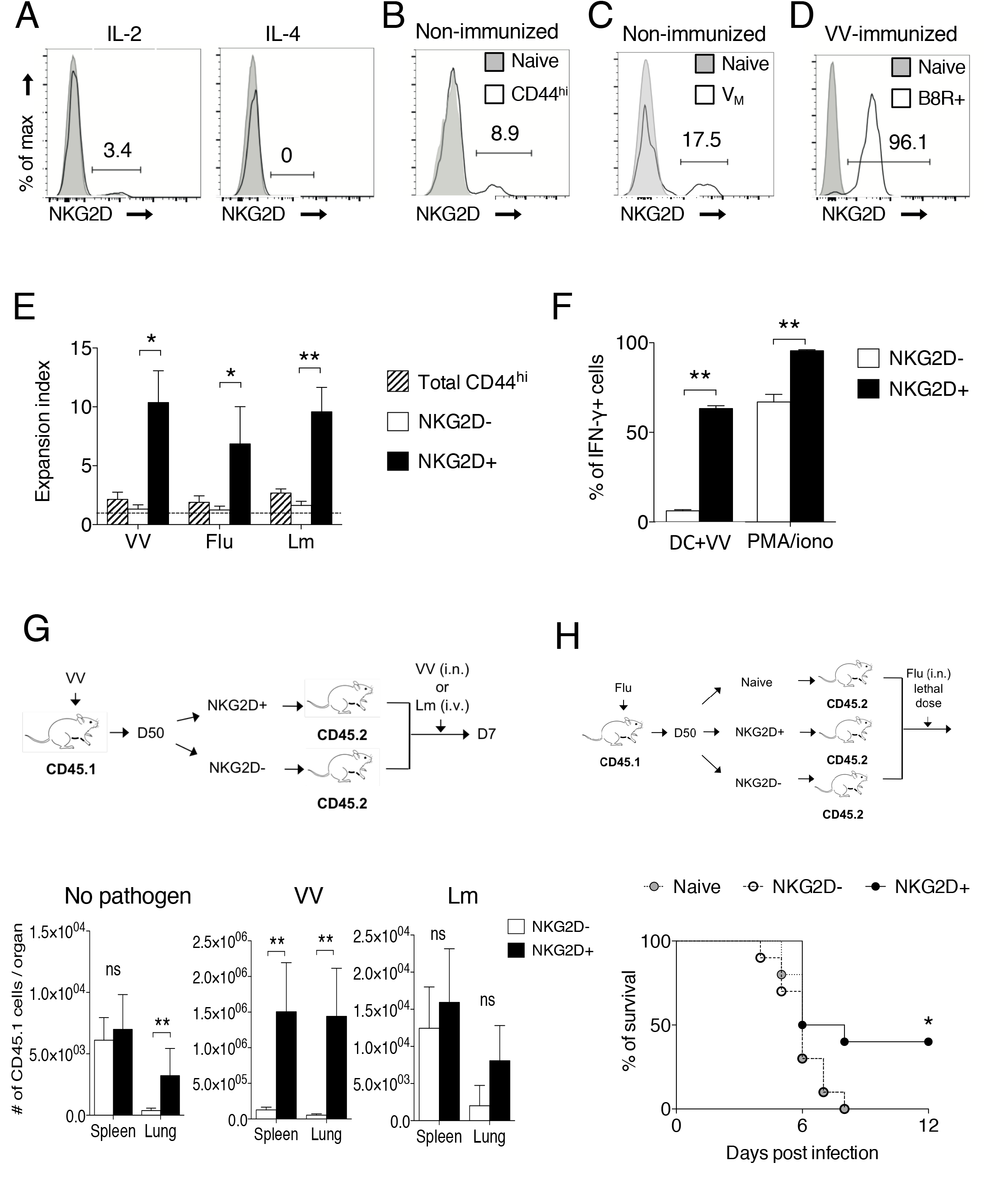
NKG2D expression identifies polyclonal antigen-induced memory CD8 T cell populations. (A) CD45.2 OTI TCR transgenic naive CD8 T cells were adoptively transferred in immunocompetent congenic mice. The next day, hosts received intra-peritoneal injections of the indicated γc cytokine antibody-complex as described in the methods. 30 days after transfer, the expression of NKG2D by OTI CD8 T cells was assessed by flow cytometry. Black histogram: OTI CD8 T cells, grey histogram: host’s CD8 T cells. The percentage of OTI cells expressing NKG2D is indicated. One representative experiment out of four is shown (n = 3 mice per experiment). (B) NKG2D expression by memory phenotype (CD44^hi^) spleen CD8 T cells from 6 weeks naive C57BL/6 mice. Black histogram: CD44^hi^ CD8 T cells, grey histogram: naive CD8 T cells. The percentage of NKG2D positive cells among CD44^hi^ CD8 T cells is shown. Results are representative of at least three experiments. (C) NKG2D expression by Virtual Memory (V_M_) CD8 T cells (B8R-specific CD44^hi^ CD49-from non-immunized naive C57BL/6 mice). Black histogram: V_M_ CD8 T cells, grey histogram: naive CD8 T cells. The percentage of V_M_ cells expressing NKG2D is indicated. Results are representative of two experiments. (D) B8R-specific memory CD8 T cells (B8R+) from vaccinia virus (VV)-immunized C57BL/6 mice (> 100 days post infection) were assessed for their expression of NKG2D (black histogram). Naive CD8 T cells from same mice were used as control. The percentage of NKG2D positive cells among B8R-specific cells is shown. Results are representative of at least three experiments. (E) C57BL/6 mice were infected with VV (intranasal), Influenza virus (Flu, intranasal) or *Listeria monocytogenes* (Lm, intravascular) and the number of total, NKG2D− and NKG2D+ CD44^hi^ CD8 T cells was measured in the blood 55 days following pathogen infection. Graph shows mean expansion index (+/− SD) of the indicated CD44^hi^ CD8 T cell populations. The dotted line represents an absence of expansion (Index = 1). One representative experiment out of two (n = at least 6 mice per group and per experiment). Wilcoxon matched-pairs signed rank test, *: pvalue < 0.05, **: pvalue < 0.01. (F) NKG2D+ and NKG2D− memory CD8 T cells were sorted from VV-infected C57BL/6 mice (55 days post infection). Cells were stimulated for 6 hours with VV-infected DC2.4 cells (DC+VV) or with PMA/ionomycin (PMA/iono) in the presence of Golgistop. The percentage of IFN-γ+ cells was measured by intracellular cytokine staining. Graph shows the mean percentage (+/− SD) of IFN-γ+ cells among each cell population. One representative experiment out of three (n = 5 mice per experiment). Mann-Withney test, **: pvalue < 0.01. (G) NKG2D+ and NKG2D− memory CD8 T cells were sorted from VV-infected CD45.1 C57BL/6 mice (50 days post infection) and 10^5^ cells were transferred in separate CD45.2 C57BL/6 congenic mice. The next day, hosts were infected with VV, Lm or left uninfected. Graphs show the mean numbers (+/− SD) of CD45.1 donor cells recovered 7 days post infection in the indicated organs. One representative experiment out of four (n = 6 mice per group and per experiment). Mann-Whitney test, ns: not significant, **: pvalue < 0.01. (H) Splenic naive, NKG2D+ and NKG2D− memory CD8 T cells were cell-sorted from Flu-infected C57BL/6 mice (45 days post infection) and 10^5^ cells were transferred in separate host mice. The next day, hosts were infected with a lethal dose of Flu. Graph shows the percentage of survival observed among each group of mice. One representative experiment out of two (n = 10 mice per group and per experiment). Log-rank (Mantel-cox) test, *: pvalue < 0.05.

To further validate NKG2D as a marker of AI memory T cells, we next analyzed NKG2D expression by VV-specific memory T cells recognizing other epitopes than B8R. Indeed, VV harbors at least 40 epitopes recognized by CD8 T cells ^30^. To extend this analysis to the whole viral epitope repertoire, memory CD8 T cells from VV-immunized mice were restimulated with VV-infected Dendritic Cells (DC). More than half of NKG2D+ memory phenotype CD8 T cells produced IFN-γ following VV restimulation, whereas only about 5% of NKG2D− memory phenotype CD8 T cells did (Figure 1F). This lack of IFN-γ production was not due to a functional defect of NKG2D− memory phenotype CD8 T cells, as restimulation with PMA and ionomycin led to IFN-γ production. As some epitopes of VV might be expressed in a delayed fashion and might not be presented in the time frame used for *in vitro* restimulation, we also performed *in vivo* rechallenge of memory CD8 T cells (Figure 1G). Although NKG2D− and NKG2D+ memory CD8 T cells from VV-infected mice equally grafted in the spleen of host mice, only NKG2D+ memory phenotype CD8 T cells had strongly proliferated, seven days after rechallenge with VV, as revealed by the number of donor cells recovered in the spleen and lung of host mice (Figure 1G). In contrast, the number of donor NKG2D− memory phenotype CD8 T cells remained close to the one observed in unimmunized host mice, revealing marginal expansion following VV rechallenge. NKG2D+ memory CD8 T cell proliferation was strictly dependent on Ag recognition, as infection with the heterologous pathogen Lm did not lead to their expansion (Figure 1G). Finally, we compared the capacity of NKG2D+ and NKG2D− memory phenotype CD8 T cells to protect naive mice against a lethal dose of virus. To do so, naive CD8 T cells as well as NKG2D− and NKG2D+ memory CD8 T cells from Flu-immune mice were transferred in naive hosts that were infected with a lethal dose of Flu. NKG2D+ memory CD8 T cells induced a significant protection of host mice as more than 40% of them survived the infection (Figure 1H). This is in contrast to naive cells and NKG2D− memory CD8 T cells that conferred no protection. Altogether, these results indicate that NKG2D expression identifies antigen-induced memory CD8 T cells in polyclonal settings, hereafter referred to as Antigen-Induced (AI) memory CD8 T cells, allowing to distinguish them from Innate memory (IN) CD8 T cells that do not express this marker.

### AI and IN memory CD8 T cells are distinguished by their TCR repertoire and cytokine secretion capacity

Next, taking advantage of NKG2D as a marker discriminating AI and IN memory CD8 T cells, we compared these two populations in terms of TCR repertoire and effector functions. A multiplex PCR identifying β chain locus VJ rearrangements showed that, as expected, AI memory CD8 T cells have a less diverse TCR repertoire than naive CD8 T cells, reflecting antigen selection (Figure 2A). In contrast, IN memory CD8 T cells have a TCR repertoire that is as diverse as naive CD8 T cells. A Principal component analysis (PCA) indicated that the TCR repertoire was more similar within the NKG2D+ subset of different mice than within the CD8 cells i.e. the naïve, NKG2D+ and NKG2D− of a given mouse (Figure 2A). As IN cells have a diverse repertoire that seems to differ in its composition from naive CD8 T cells, we tested if these cells could participate to a primary immune response against a pathogen, i.e. whether they can mount responses against unknown foreign epitopes. Equal numbers of CD45.1 naive and CD45.2 IN memory CD8 T cells were sorted from naive mice and co-transferred to congenic CD45.1/CD45.2 hosts that were infected the next day with VV or Lm. Seven days post infection, the contribution of transferred cell populations to the primary response was determined in the spleen (Figure 2B). Both naive and IN memory cells expanded during the primary immune response and became NKG2D positive. We also evaluated the relative contribution of the two grafted cell populations to the NKG2D+ CD44^hi^ effector cells response. During VV infection, naive and IN memory CD8 T cells generated almost equal numbers of NKG2D+ CD44^hi^ effector cells. In contrast, during Lm infection, IN memory CD8 T cells generated more NKG2D+ CD44^hi^ effector cells compared to naive cells (Figure 2B). In conclusion, IN cells have a more diversified repertoire than AI and can contribute to a primary T cell response.

**Figure 2:**
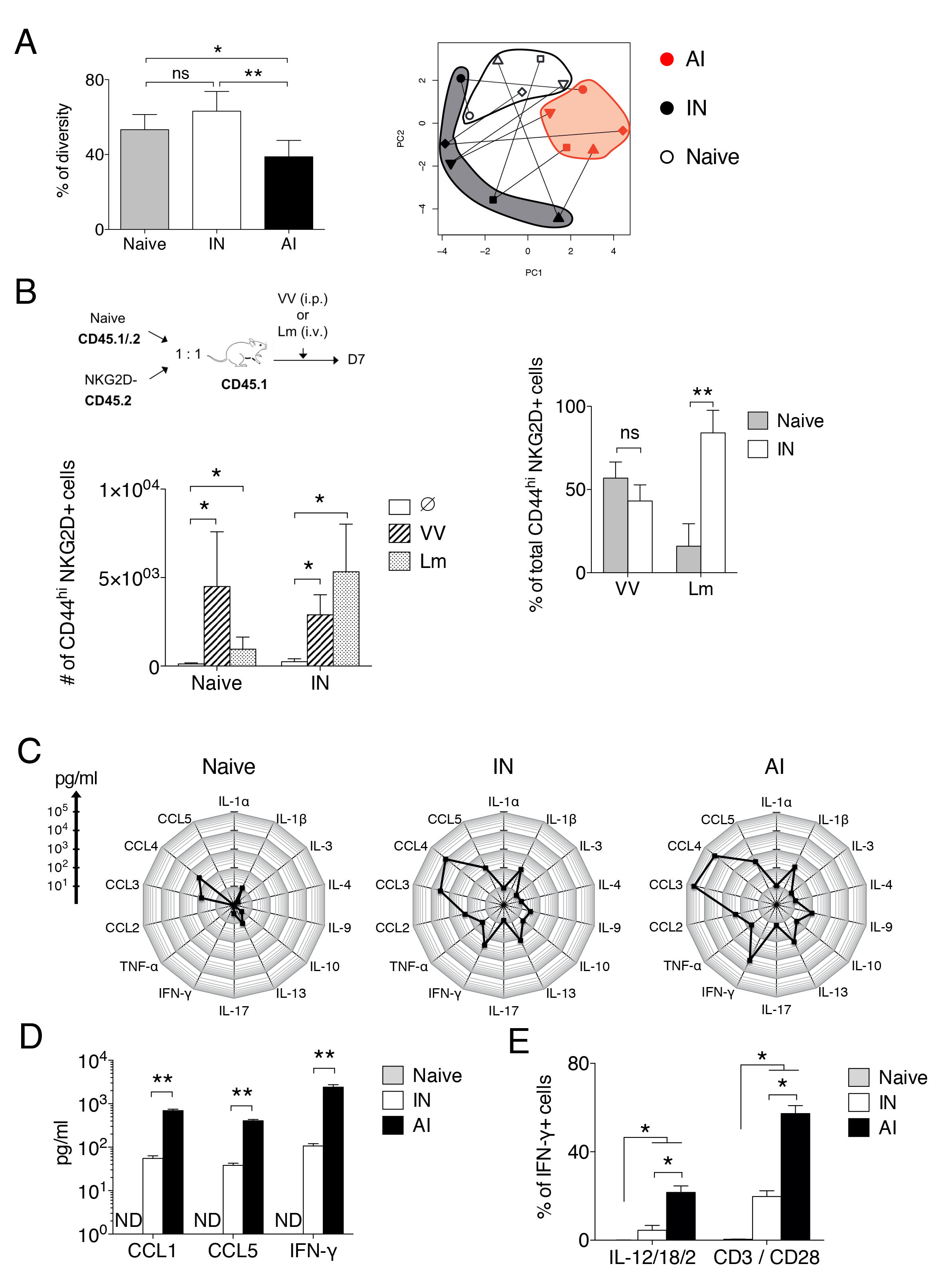
Characterization of IN memory CD8 T cells. (A) Naive, IN and AI memory CD8 T cells were sorted from VV-infected C57BL/6 mice (50 days post infection) and multiplex PCR was performed on genomic DNA to detect VβJβ rearrangements among each cell population. Left graph shows the mean percentage (+/− SD) of TCR repertoire diversity, calculated as described in the methods. Mann-Whitney test (n = 5 mice), ns: not significant, *: pvalue < 0.05, **: pvalue < 0.01. Right: Principal component analysis based on the presence or absence of each possible VβJβ rearrangement among each cell population. This representation shows the distribution of CD8 T cell populations regarding to the composition of their VβJβ TCR repertoire. Each form of symbol represents a mouse and lines connect cell populations sorted from the same mouse. (B) Naive and IN memory CD8 T cells were sorted from naive CD45.1/CD45.2 and CD45.2 C57BL/6 mice respectively and were co-transferred at a 1:1 ratio in congenic CD45.1 C57BL/6 mice. Host mice were infected with VV, Lm or left uninfected (Ø). Left graph shows the mean number (+/− SD) of CD44^hi^ NKG2D+ CD8 T cells in the spleen generated from transferred cell populations 7 days post infection. Right graph shows the contribution of each transferred CD8 T cell population to the total CD44^hi^ NKG2D+ CD8 T cells generated from transferred cells (mean percentage +/− SD). One representative experiment out of two (n = 5 mice per group and per experiment). Mann Whitney test, ns: not significant, **: pvalue < 0.01. (C) Equal number of naive, IN and AI memory CD8 T cells were sorted from VV-infected mice (> 100 days after infection) and were restimulated for 12 hours with anti-CD3 plus anti-CD28 antibodies in the presence of IL-2. Radar plots show the mean amount of each cytokine produced, measured by multiplex. One experiment (n = 5 mice). (D) Graph shows the mean amount (+/− SD) of memory-associated poised cytokines produced by CD8 T cell populations, measured by ELISA. One representative experiment out of three (n = 5 mice per experiment). Mann-Whitney test, **: pvalue < 0.01. ND: not detected. (E) Production of IFN-γ by naive, IN and AI memory CD8 T cells from VV-infected mice (50 days post infection) was measured by intracellular cytokine staining following 5 hours of restimulation with anti-CD3 plus anti-CD28 antibodies or with a mixture of IL-12/IL-18/IL-2. Graph shows the mean percentage (+/− SD) of IFN-γ+ cells among each CD8 T cell population. A pool of two representative experiments is shown (n = 4 mice in total). Mann-Whitney test, *: pvalue < 0.05.

One key property of memory CD8 T cells is the rapid and increased production of cytokines and chemokines upon TCR stimulation. We therefore compared the capacity of polyclonal IN and AI memory and naive CD8 T cells to produce cytokines and chemokines in response to TCR stimulation. Early after TCR triggering, both IN and AI produced a broader array of cytokines/chemokines than naive cells. These factors were also secreted in larger amounts by memory T cells of both subsets (Figure 2C). Importantly, the same pattern of cytokines/chemokines was produced by IN and AI memory cells, although AI produced at least a tenfold higher quantity of the poised memory cytokine/chemokines CCL5, CCL1 and IFN-γ than IN (Figure 2D). AI memory cells can also produce IFN-γ upon stimulation by innate signals such as IL-12 and IL-18, a property known as innate function of memory CD8 T cells ^31^. A fraction of IN memory cells produced IFN-γ upon IL-12/IL-18 stimulation, albeit at a lower frequency than AI memory CD8 T cells (Figure 2E). Taken together, these results indicate that although AI and IN memory CD8 T cells produced a similar pattern of cytokines following stimulation, AI produce higher amounts of cytokines than IN memory T cells.

### Transcriptome analysis of AI and IN memory CD8 T cells

To further characterize the two subsets defined by NKG2D expression, we compared their transcriptome using microarrays. F5 TCR transgenic CD8 T cells were transferred in naive mice before infection with VV-NP68, to establish an internal control of AI memory CD8 T cells. Eighty days post infection, F5 memory cells as well as host’s IN and AI memory CD8 T cell populations were sorted and their transcriptome was analyzed. Naive CD8 T cells were sorted from naive F5 and C57BL/6 mice. We first performed a PCA analysis that shows that 70% of the variability of the samples is explained by the first two principal components (PC). Samples were aligned along PC1 according to their differentiation stage, while PC2 highlighted a difference between monoclonal F5 TCR transgenic T cells and polyclonal CD8 T cells (Figure 3A). F5 and polyclonal CD8 T cells differed by the expression of few genes, among which those coding for the TCR, reflecting the monoclonality of the F5 CD8 population (Figure S5A) and one set of genes encoding the inhibitory NK cell receptors (*Klra*) was expressed by polyclonal AI memory cells but not F5 memory cells. Accordingly, these Ly49 receptors are exclusively expressed by a small fraction of polyclonal memory CD8 T cells in contrast to TCR transgenic F5 memory cells (Figure S5B). This property is shared by both IN and AI polyclonal memory cells (Figure S5C). Importantly, all genes differentially expressed by AI memory CD8 T cells compared to naive cells were also differentially expressed by F5 CD8 T cells when comparing memory to naive cells confirming their antigen-induced nature (data not shown). We then compared AI and IN polyclonal memory populations. IN memory cells were positioned closer to AI memory cells than to naive CD8 T cells on the PC1 axis, confirming their memory differentiation (Figure 3A). Indeed, AI and IN memory CD8 T cell transcriptomes differ in the expression levels of several genes encoding transcription factors, effector molecules, NK cell receptors, chemokine receptors and integrins (Figure 3B). Compared to IN, AI memory CD8 T cells express higher levels of genes encoding effector molecules involved in the killing of target cells through cytotoxicity. Accordingly, this memory cell population displays a higher capacity to mediate killing, in anti-CD3 redirected cytotoxicity assay (Figure S6). Importantly, AI memory CD8 T cells express higher levels than IN cells of transcription factors that promote the full differentiation of memory CD8 T cells, such as Tbet, ID2, Zeb2 and Blimp-1 (Figure 3C and 3D). In contrast, IN memory CD8 T cells show increased levels of transcription factors that promote a less differentiated state of memory CD8 T cells, such as Eomes and ID3. The pattern of transcription factors expressed by AI memory cells is thus in accordance with their more differentiated state compared to IN.

**Figure 3:**
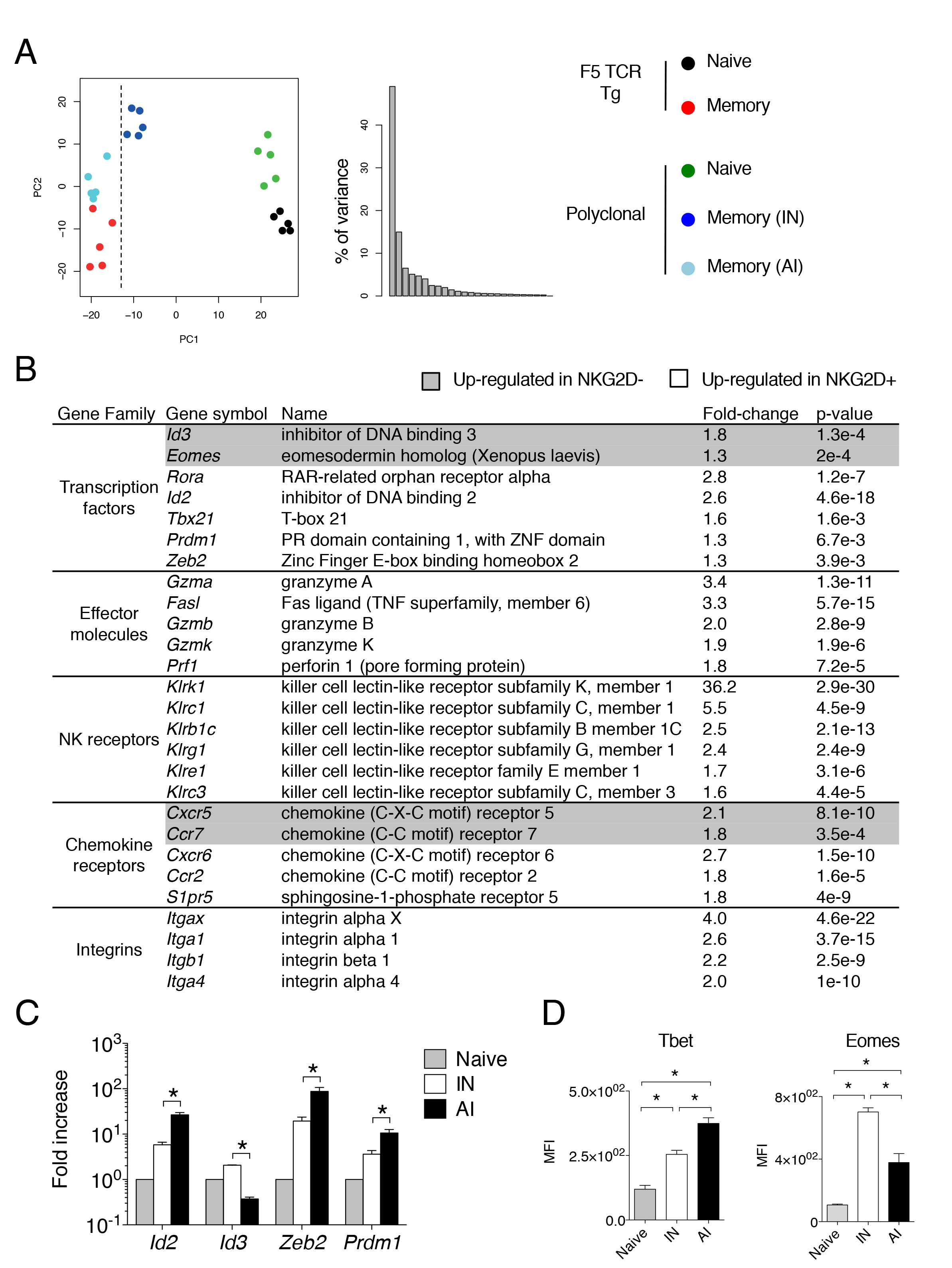
Transcriptome analysis of AI and IN memory CD8 T cells. Naive CD45.1 F5 TCR transgenic CD8 T cells were adoptively transferred in congenic host mice that were subsequently immunized with VV-NP68. 80 days post infection, F5 (TCR Tg) memory cells and polyclonal NKG2D− and NKG2D+ memory CD8 T cells from the host were sorted from 5 independent groups of 8 mice. As a control, naive F5 and polyclonal CD8 T cells were sorted from 5 independent groups of 3 naive F5 and C57BL/6 mice respectively. The transcriptome of these cell populations was compared by microarrays. (A) Principal component analysis was performed on whole microarray data. Left graph shows the distribution of samples according to Principal Components 1 and 2 (PC1 and PC2). Right graph shows the percentage of variance explained by successive principal components. (B) The main genes differently expressed between polyclonal IN and AI memory CD8 T cells are listed. Fold changes and p-values are indicated for each gene. The family to which each group of genes belongs is also indicated. (C) Naive, IN and AI memory CD8 T cells were sorted from VV-infected mice (50 days post infection). The expression level of several transcription factors by each cell population was assessed by quantitative PCR. Graph shows the mean fold increase (+/− SD) compare to naive CD8 T cells. One representative experiment out of two (n = 5 mice per experiment). (D) The expression of Tbet and Eomes by naive, IN and AI memory CD8 T cells from VV-infected mice (50 days post infection) was assessed by flow cytometry. Graphs show mean of Mean Fluorescence Intensity (+/− SD) of each transcription factors. One representative experiment out of two (n = 4 mice per experiment). Mann-Withney test, *: pvalue < 0.05.

### AI memory CD8 T cells have an increased capacity to enter inflamed peripheral tissues compared to IN memory cells

One important characteristic of memory cells is their capacity to circulate and migrate to inflamed tissues ^32–34^. As IN cells express less memory-specific chemokine receptors and integrins than AI cells, we compared their ability to traffic to the lung upon inflammation or infection. Inflammation was first induced in mouse lungs by the TLR3 agonist poly(I:C) that induces the production of type-I IFN and its downstream chemokines (CXCL9, 10 and 11) ^35^. Memory CD8 T cells containing similar numbers of IN and AI populations were purified from VV-infected mice and transferred into host mice that then received intranasal injection of poly(I:C). The recruitment of AI and IN memory cells in different organs was assessed two days after poly(I:C) injection. The same numbers of donor AI and IN memory CD8 T cells were found in the spleen and in the blood of recipient mice injected with PBS or poly(I:C) (Figure S7) as reflected by a cell number ratio close to 1 (Figure 4A). As expected the recrutement of donor memory CD8 T cells within the lung parenchyma and airways was induced following poly(I:C) injection (Figure S7). However, AI memory CD8 T cells were preferentially recruited compared to IN cells (Figure 4A). Following lung infection by a pathogen spleen memory cells are recruited to the lung in an antigen independent fashion ^36 37^. To analyse the capacity of AI and IN memory CD8 T to be attracted to the infected lung, VV-specific memory CD8 T cells containing NKG2D− and NKG2D+ populations were isolated from VV-immune mice and transferred into host mice that had been immunized two days before with Flu virus and the recruitment was assessed two days later. Again, AI memory CD8 T cells were preferentially recruted to the lung during Flu infection, accumulating within the parenchyma (Figure 4B). In conclusion, our results show that upon inflammation or infection, AI memory CD8 T cells enter the lung parenchyma more efficiently than IN memory CD8 T cells.

**Figure 4:**
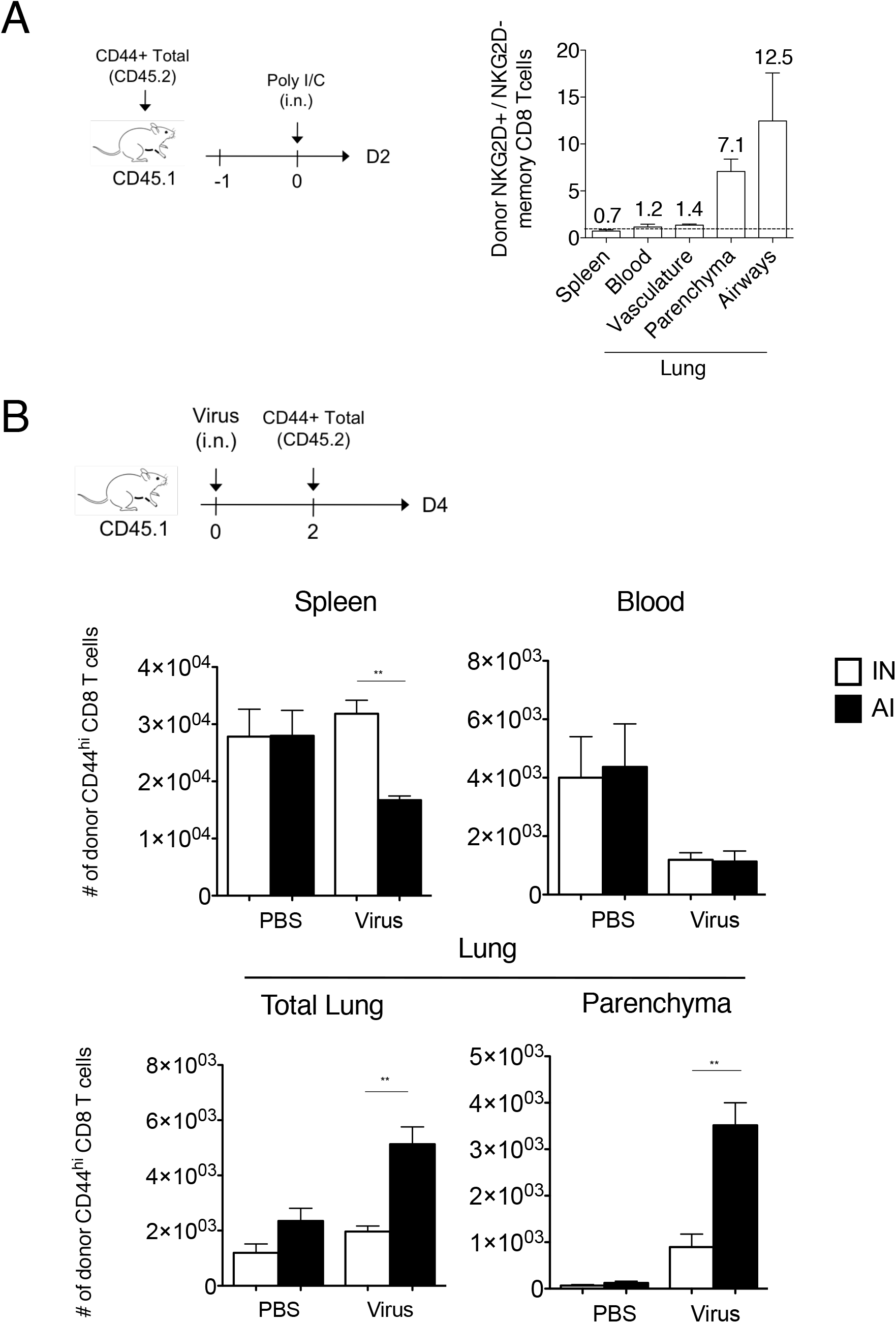
IN have a reduced capacity to access inflamed peripheral tissues compared to AI memory CD8 T cells. (A) CD45.2 memory CD8 T cells were purified from VV-infected mice (intraperitoneal immunization) and transferred into congenic CD45.1 mice. The next day, host mice received intranasal administration of poly(I:C) or PBS. Two days later, the numbers of donor IN and AI memory CD8 T cells were measured in various organs. Graph shows the mean ratios (+/− SD) between donor IN and AI memory CD8 T cells in spleen, blood and lung. Lung intravascular staining was performed to differentiate cells in the vasculature and in the parenchyma. One representative experiment out of two (n = 5 mice per group per experiment). (B) CD45.2 memory CD8 T cells were purified from VV-infected mice (intraperitoneal immunization) and transferred into congenic CD45.1 mice immunized with Flu virus (intranasal immunization) two days earlier. Graphs show mean numbers (+/− SD) of donor IN and AI memory CD8 T cells in the spleen, the blood and the lung (total or in the parenchyma) two days later. Mann-Whitney test, **: pvalue < 0.01.

### Recruitment of AI memory CD8 T cells into the inflamed lung is CXCR3 and CD49a/CD49d dependent

Memory CD8 T cell trafficking to the lung parenchyma and airways has been reported to be dependent on ITGA1-4 integrins and chemokine receptors CXCR3 ^32^ ^34 38^. Using CXCR3 KO mice we confirmed that CXCR3 expression is required for AI memory cells recrutement to the lung parenchyma (Figure 5A). However, the differential recruitment of AI and IN memory subsets did not involve CXCR3 as both memory cell types expressed similar level of CXCR3 and migrated strongly towards CXCL10 in a transwell assay (Figure 5B). By contrast, AI differ from IN by the expression level of several integrin mRNA namely Itga1 and Itga4 and Itgb1 encoding for the alpha and beta chains of VLA1 (CD49a, CD29) and VLA4 (CD49d, CD29), respectively (Figure 3B, ^26 39^), a difference confirmed at the protein level by flow cytometry analysis (Figure 5C). We thus tested the if antibodies directed against CD49a and CD49d could alter the recruitment of AI to the infected lung. Antibody treatment did not affect AI numbers in spleen and blood (Figure S8). In contrast, it significantly inhibited AI recruitment to the lung parenchyma (Figure 5D). Altogether, these results indicate that following lung infection, AI memory cells are the main memory subset recruited to the lung parenchyma, a process that is dependent on CXCR3 and CD49a/CD49d integrins expression.

**Figure 5:**
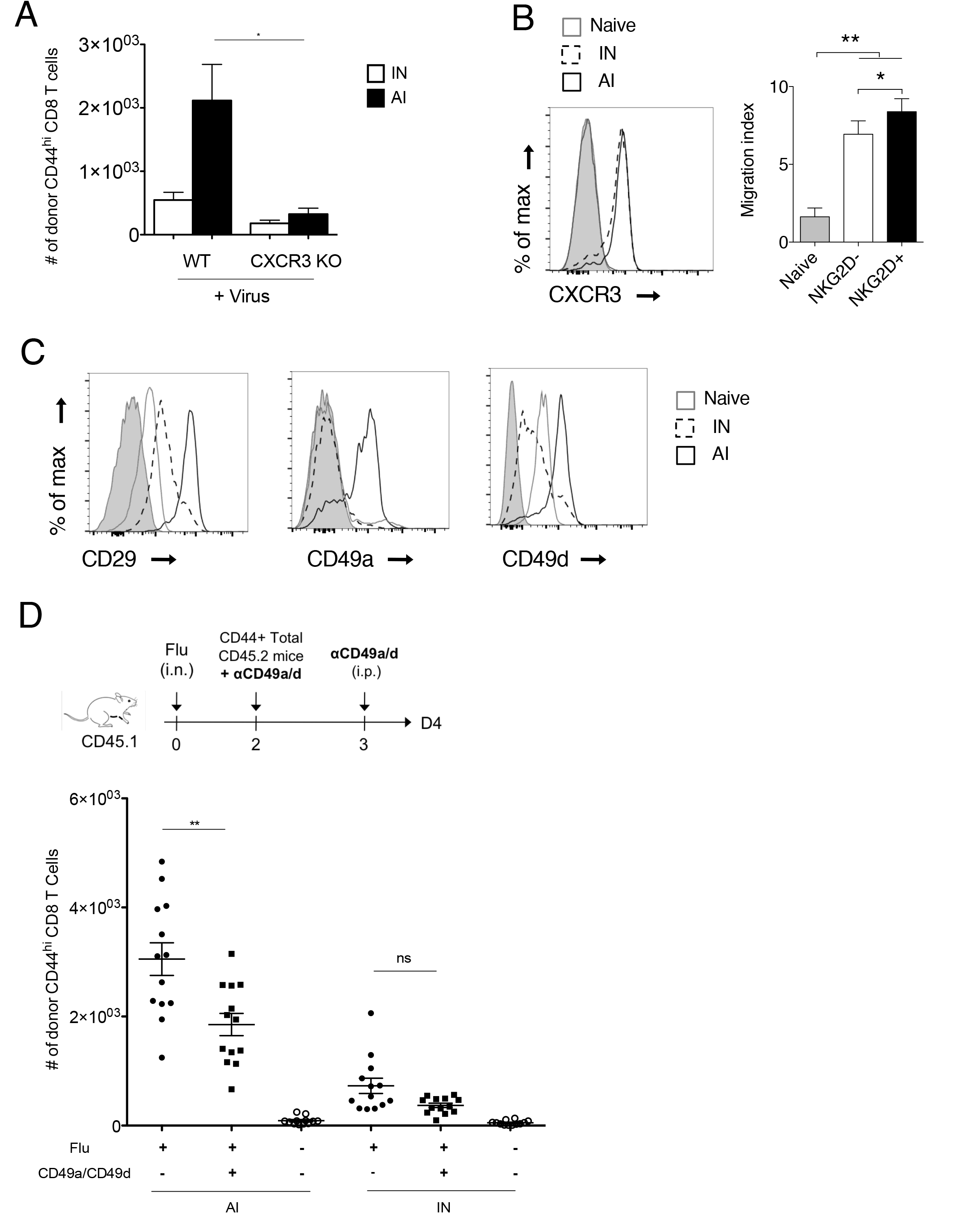
Role of CXCR3 and CD49a/CD49d for entry of AI memory CD8 T cells into inflamed lung. (A) Memory CD8 T cells purified from VV-infected mice C57BL/6 mice or CXCR3 KO mice (intraperitoneal immunization) were transferred into congenic hosts. 35 days later, host mice received intranasal administration of Flu virus. Two days later, the number of donor IN and AI memory CD8 T cells was measured in lung parenchyma. One representative experiment out of two. Mann-Whitney test, *: pvalue < 0.05. (B) Histogram shows the expression of CXCR3 by naive, IN and AI memory CD8 T cells from VV-infected mice (50 days post infection). Grey histogram: control isotype. Graph shows the migration of naive, IN and AI memory CD8 T cells from VV-infected mice toward CXCL10 in a transwell assay (mean +/− SD). One representative experiment out of three (n = 5 mice per experiment). Mann-Whitney test, *: pvalue < 0.05, **: pvalue < 0.01. (C) The expression of CD29, CD49a and CD49d by naive, IN and AI memory CD8 T cells from VV-infected mice (50 days post infection) was assessed by flow cytometry. Grey histogram: control isotype. One representative experiment out of three (n = 2 mice per experiment). (D) Memory CD8 T cells were purified from VV-infected mice (intranasal immunization), incubated or not with 1 μg/ml anti-CD49a and anti-CD49d antibodies, and then transferred into congenic mice that had been immunized intranasaly 2 days before with Flu virus. The next day, host mice received intraperitoneal administration of 250 μg/ml anti-CD49a plus anti-CD49d antibodies or PBS. Two days later, the number of donor IN and AI memory CD8 T cells was measured in lung parenchyma. Graph shows mean numbers (+/− SD) of donor IN and AI memory CD8 T cells. A pool of two representative experiments is shown (n = 13 mice in total). Mann-Whitney test, *: pvalue < 0.05, **: pvalue < 0.01.

## Discussion

In this study, we demonstrated that NKG2D surface expression is restricted to AI memory CD8 T cells, allowing their discrimination from IN memory CD8 T cells generated under physiological conditions. We showed that this dichotomy is conserved in different mouse strains and in response to infection by different pathogens. Expression of NKG2D by virtual memory CD8 T cells has recently been reported at the mRNA level ^39^. We found that sorted NKG2D−negative memory-phenotype IN cells containing the virtual memory cells expressed higher levels of mRNA encoding NKG2D compared to naive cells (FC=2, Figure S9). However, we found that the small fraction of B8R+ virtual memory (0.02%) within CD44^hi^ CD49d negative CD8 T cells found in naive mice are mainly NKG2D negative at the protein level (Figure 1C). Importantly, NKG2D positive AI memory CD8 T cells expressed much higher levels of NKG2D mRNA than IN in quiescent cells which could explain the difference observed in NKG2D protein levels. Thus, the surface expression of NKG2D protein is restricted to AI memory CD8 T cells.

Taking advantage of NKG2D expression, we compared IN memory CD8 T cells to pathogen-induced memory CD8 T cells. We performed a transcriptome comparison of IN and AI memory CD8 T cell populations. Our results clearly demonstrated that VV-induced memory cells whether polyclonal (NKG2D+ CD8 T cells) or monoclonal (F5 TCR transgenic CD8 T cells) share the same transcriptome. IN have also acquired a genetic program typical of memory, nevertheless, these cells are less differentiated compared to AI memory CD8 T cells. In agreement with a previous study ^40^, IN did not express a specific gene expression pattern that could indicate an independent differentiation pathway. This suggests that IN memory CD8 T cells represent an intermediate stage of differentiation between naive and AI memory CD8 T cells rather than a distinct CD8 T cell lineage. This is also supported by their cytokine secretion pattern in response to TCR stimulation similar to that of AI memory CD8 T cells. Differentiation of memory cells is regulated by different pairs of transcription factors, such as Tbet and Eomes, Blimp1/Bcl6 or ID2/ID3 ^41^. Interestingly, genes encoding for these transcription factors are differentially expressed between IN and AI memory CD8 T cells. Indeed, AI memory CD8 T cells express higher levels of genes encoding for transcription factors that promote memory CD8 T cell full differentiation (*Tbx21, Id2, Prdm1, Zeb2*). In contrast, IN memory CD8 T cells express higher levels of genes encoding transcription factors that favor a less differentiated state (*Id3, Eomes*). Thus, the expression pattern of transcription factors observed in IN and AI memory CD8 T cell populations fits with the observed differentiation state. In agreement with their transcription factors expression pattern, AI memory CD8 T cells express higher levels of genes encoding for effector molecules, such as granzymes, perforin and Fas ligand, confirming that these cells are more differentiated and are able to kill a potential target more rapidly.

One major difference between naive and memory cells is the capacity of memory cells to access and reside within tissues parenchyma. Following a pulmonary infection, pathogen-specific CD8 effector cells enter the lung parenchyma and under specific signaling some of them differentiate in Trm that ensure a robust response if a secondary infection occurs ^3 4^. In the lung, in contrast to other tissues, the population of Trm wanes over time ^5 6^. Thus, long term protection of the lung relies on the recruitment of secondary memory cells stored in lymphoid tissues. Indeed, following infection of the lung, inflammation rapidly induces the recruitment of memory cells independently of their antigenic specificity ^36,37,42^, although pathogen specific spleen memory T cells also rapidly gain access to the tissue ^2^.

Previous studies have demonstrated that the CD49d integrin is differentially expressed between B8R-specific IN and AI memory CD8 T cells ^26,22^. Transcriptome comparison of IN and AI revealed here that other integrin chains are also differentially expressed by AI and IN memory cells. This was confirmed at the protein level: AI memory CD8 T cells expressed higher levels of several integrins (CD29, CD49a, CD49d) compared to naive or IN memory CD8 T cells. These integrins play a key role in immune cell migration, allowing them to exit blood vessels and access peripheral tissues ^38^. Indeed, ITGA1/B2 (CD29, CD49a) and ITGA4/B2 (CD29, CD49d) play an essential role in the extravasation of CD8 T cells in the lung or the brain, respectively ^43,44^. Accordingly, we found that upon lung inflammation, AI memory CD8 T cells were preferentially recruited within the lung parenchyma and we demonstrated that this process was dependent on integrins. In contrast, IN memory CD8 T cells remained within the lung vasculature. Memory CD8 T cell trafficking to the lung parenchyma and airways has been reported to be dependent of the chemokine receptor CXCR3 ^34^. In agreement, CXCR3 KO AI memory cells were not recruited to the lung. Importantly, we did not observe any differential CXCR3-expression or CXCL10-induced migration between IN and AI memory CD8 T cells. This indicates that CXCR3 expression is necessary but not sufficient to access the inflamed lung parenchyma.

IN memory CD8 T cells have a TCR repertoire as diverse as that of naive CD8 T cells, although their repertoire does not completely overlap. This diversified TCR repertoire could allow IN memory CD8 T cells to contribute to multiple immune responses. In line with this, we showed that IN memory CD8 T cells participated to primary immune responses against two pathogens, namely VV and Lm. Participation of IN memory CD8 T cells to primary immune responses against infectious pathogens could significantly increase the efficiency of these responses. Indeed, we showed that physiologically generated IN produced a lot more cytokines than naive cells when stimulated through their TCR. In line with this, Lee and colleagues demonstrated that OVA-specific IN memory CD8 T cells cleared Lm-OVA infection more efficiently than naive CD8 T cell do ^27^. Similarly, results obtained in mice that are deficient in IL-4-induced IN indicate a decreased capacity to control a primary LCMV infection in the absence of these cells ^45^. The protection conferred by IN cells could be direct, as a result of their enhanced effector functions or indirect, through the help to naive CD8 T cells ^46^. However, our results show that they are not recruited to the inflamed lung parenchyma due to the lack of ITGA1/4 integrins expression. This is in line with the demonstration that virtual memory CD8 T cells do not confer protection against *Listeria monocytogenes* upon gut infection ^39^.

The decreased capacity of IN memory cells to access non-lymphoid tissues in response to inflammatory chemokines could be essential for the prevention of autoimmunity. Due to its generation process, the population of IN memory CD8 T cells might be preferentially generated from naive cells with increased sensitivity for self-antigens. Indeed, CD5^hi^ naive CD8 T cells, that have an increased sensitivity to self antigens, are more prone to undergo LIP compared to CD5^lo^ naive cells ^47 48^. Moreover, CD5^hi^ naive cells are more predisposed to become virual memory CD8 T cells compared to CD5^lo^ naive cells. Of note, increased lymphopenia drives the development of auto-aggressive T cells in NOD mice. This mechanism accounts partly for the development of diabetes in this mouse model ^49^. The exclusion from peripheral tissues of IN memory CD8 T cells that show increased reactivity compared to naive cells might thus be important to avoid autoimmunity.

In conclusion, NKG2D is a novel marker of AI memory cells. Moreover, although AI and IN memory CD8 T cells are similar from a phenotypic, transcriptomic and functional point of view, they differ in their capacity to be recruited to inflamed lung parenchyma in an integrin-dependent fashion.

## Material and methods

### Mice

F5 TCR (B6/J-Tg(CD2-TcraF5,CD2-TcrbF5)1Kio/Jmar) transgenic mice were provided by Professor D. Kioussis (National Institute of Medical Research, London, U.K.) and backcrossed on CD45.1 C57BL/6 background ^50^. The F5 TCR recognizes the NP68 peptide from influenza A virus (ASNENMDAM) in the context of H2-D^b^. OTI TCR transgenic (B6/J-Tg(Tcra,Tcrb)1100Mjb/Crl), CD45.2 (C57BL/6J) and CD45.1 (B6.SJL-Ptprc^a^Pepc^b^/BoyCrl) C57BL/6J, BALB/c and OF1 mice were purchased from Charles River Laboratories (L’arbresle, France). The OTI TCR recognizes the OVA257-264 peptide from chicken ovalbumin (SIINFEKL) in the context of H2-K^b^. Mice were bred or housed under SPF conditions in our animal facility (AniRA-PBES, Lyon, France). All experiments were approved by our local ethics committee (CECCAPP, Lyon, France) and accreditations have been obtained from governmental agencies.

### Pathogens and mouse immunization

The recombinant influenza virus strain WSN encoding the NP68 epitope (Flu-NP68) was produced by reverse genetics from the A/WSN/33 H1N1 strain. The recombinant vaccinia virus, expressing the NP68 epitope (VV-NP68), was engineered from the Western Reserve strain by Dr. Denise Yu-Lin Teoh, in Pr. Sir Andrew McMichael’s laboratory at the MRC (Human Immunology Unit, Institute of Molecular Medicine, Oxford, UK). The *Listeria monocytogenes* strain 10403s (Lm) was produced from clones grown from organs of infected mice. For immunization, anesthetized mice received intranasal (i.n.) administration of Flu (2 × 10^5^ TCID_50_), VV (2 × 10^5^ PFU) or poly(I:C) (30 μg) in 20 μL of phosphate-buffered saline (PBS) or intravenous (i.v.) Lm (2 × 10^3^) administration in 200 μl PBS. For some indicated experiments, mice received intraperitoneal (i.p.) administration of 1 × 10^6^ PFU VV in 200 μl PBS.

### TCR transgenic memory CD8 T cells

To generate AI TCR transgenic memory CD8 T cells, 2 × 10^5^ naive CD45.1 F5 CD8 T cells were transferred in C57BL/6 mice by i.v. injection. The next day, mice were infected with VV-NP68 as described above. To generate IN TCR transgenic memory CD8 T cells, 1 × 10^6^ naive OTI CD8 T cells were transferred in sub-lethally irradiated (600 rad) CD45.1 C57BL/6 mice by i.v. injection. OTI naive cells were also transferred to immunocompetent mice that further received i.p. injections of 1.5 μg of IL-2 or IL-4 (Peprotech) immunocomplexed to anti-IL-2 (S4B6, BioXcell) or anti-IL-4 (11B11, BioXcell) antibody, during 7 consecutive days.

### Cell preparation and flow cytometry

To discriminate between tissue resident and circulating memory CD8 T cells, *in vivo* intravascular staining was performed as previously described ^51^. Briefly, mice were injected i.v. with 3 μg of CD45-BV421 antibody (BioLegend) diluted in 200 μl of sterile PBS (Life Technologies) and were sacrificed 2 min after injection by overdose of pentobarbital. Blood samples (100 μl) were collected on EDTA by retro-orbital bleeding. Spleen and lymph nodes were harvested, mechanically disrupted and filtered through a sterile 100 μm nylon mesh filter (BD). To collect broncho-alveolar lavages (BAL), the trachea was exposed and cannulated with a 24-gauge plastic catheter (BD Biosciences) and lung were lavaged twice with 1 mL cold sterile PBS. Lungs were enzymatically digested using a specific dissociation kit and following manufacturer’s instructions (Miltenyi Biotec).

Surface staining was performed on single cell suspensions from each organs for 30 min at 4°C with the appropriate mixture of monoclonal antibodies diluted in staining buffer (PBS supplemented with 1% FCS (Life Technologies) and 0.09% NaN3 (Sigma-Aldrich, Saint Quentin-Fallavier, France)). To identify B8R-specific memory CD8 T cells, dextramer staining was performed 20 min at room temperature using B8R dextramer (Immudex), before surface staining. The following antibodies (clones) were used for surface staining: NKG2D (CX5), CD45.1 (A20), CD45.2 (104), CD122 (TM-b1), CD62L (MEL-14), CD8 (53-6.7), CD44 (IM7.8.1), CXCR3 (Cxcr3-173), CD49a (HA 31-8), CD49d (R1-2), CD29 (eBioHMb1-1), CD11c (N418). To perform intracellular cytokine staining, cells were fixed and permeabilized using CytoFix/CytoPerm (BD Pharmingen). To detect transcription factors, FoxP3 Kit (eBioscience) was used to fix and permeabilize cells. The following antibodies (clones) were used for intracellular staining: IFN-γ (XMG1.2), CCL5 (2E9), Tbet (4B10) and Eomes (Dan11mag). All analyses were performed on a Becton Dickinson FACS LSR II or Fortessa and analysed with FlowJo software (TreeStar, Ashland, OR, USA).

### In vitro stimulation and cytokines production measurements

For measurements of cytokine production, 5×10^4^ naive and memory (NKG2D− and NKG2D+) CD8 T cells sorted from VV-infected mice were cultured for 12 hours with plate bound anti-CD3 antibody (145-2C11, 10 μg/mL, BD Biosciences), soluble anti-CD28 antibody (37.51, 1 μg/mL, BD Biosciences) and IL-2 (2%). Supernatants were collected and cytokine production was measured by bead-based multiplexing technology for IL-1α, IL1-β, IL-3, IL-4, IL-9, IL-10, IL-13, IL-17, IFN-γ, TNF-α, CCL2/3/4/5 (Bio-Plex Pro, Bio-Rad) or by ELISA for CCL1, CCL5, and IFN-γ (Mouse DuoSet, R&D system). For flow cytometry measurements of cytokine production at single cell level, 1×10^5^ NKG2D− and NKG2D+ memory CD8 T cells, sorted from VV-infected mice, were cultured for 6 hours with VV-infected (MOI = 10) DC2.4 cells or with PMA (20 ng/mL) and Ionomycin (1 μg/mL) in the presence of golgistop (BD). Alternatively, 1×10^6^ total splenocytes from VV-infected mice were cultured for 5 hours with plate bound anti-CD3 antibody and soluble anti-CD28 antibody or with IL-12 (10 ng/mL, R&D system), IL-18 (10 ng/mL, MBL) and IL-2 (10 ng/mL, Peprotech).

### Protection assay

To evaluate the degree of protection associated with each CD8 T cells population, mice were transferred with 1×10^5^ naive, NKG2D− or NKG2D+ memory CD8 T cells from Flu-infected mice. The next day, host mice were infected with a lethal dose (1×10^6^ TCID_50_) of Flu. Mice weight loss was measured each day, for up to 12 days, after infection. Mice that lost more than 20% of initial body weight were euthanized.

### TCR repertoire analysis

Naive, NKG2D− and NKG2D+ memory CD8 T cells were sorted from VV-infected mice (50 days post infection). Cells were lyzed and multiplex PCR were performed by ImmunID (Grenoble, France) on genomic DNA to detect VJ rearrangements at the TCR β chain locus. For each cell population, the percentage of TCR repertoire diversity was calculated as the ratio between the number of observed VJ recombinations and the theoretical number of VJ recombinations, i.e. 209.

### Transcriptome analysis

F5 TCR transgenic memory CD8 T cells were generated as described above. 80 days after VV-NP68 infection, CD45.1 F5 memory CD8 T cells as well as host’s NKG2D− and NKG2D+ memory CD8 T cells were sorted from 5 pools of spleens, each from eight mice (purity > 98%). Naive F5 and polyclonal CD8 T cells were sorted from pools of three spleens from naive F5 and C57BL/6 mice respectively. Total RNA was extracted from dry cell pellets according to the “Purification of total RNA from animal and human cells” protocol of the RNeasy Micro Kit (QIAGEN, Hilden, Germany). Purity and integrity of the RNA was assessed on the Agilent 2100 Bioanalyzer (Agilent, Palo Alto, CA, USA). Total RNA from each sample was amplified, labeled and hybridized to mouse GeneChip HT MG-430 PM Plates as described in the Affymetrix GeneChip 3' IVT PLUS Reagent Kit User Manual (Affymetrix, Inc., Santa Clara, CA, USA). Affymetrix CEL files were analyzed in R using the appropriate packages from the Bioconductor suite (<u>https://www.bioconductor.org/</u>). Raw probe signals were background corrected using the maximum likelihood estimation of the normal-exponential mixture model ^52^, normalized using the variance stabilization normalization ^53^ followed by a quantile normalization ^54^. Summarization was performed using the median-^55^ using a modified version 17.1 of the Entrez-Gene based re-annotated chip description file {Dai:2005ge}. Non-informative genes were filtered using the I/NI algorithm ^56^. Linear models were applied using the limma package in order to compute the average expression level for each cell type. A random effect was introduced to account for the paired design. Statistical contrasts were then applied to compute differential expression between the different cell types. The empirical Bayes method was used to compute moderated p-values that were then corrected for multiple comparisons using the Benjamini and Hochberg's false discovery rate (FDR) controlling procedure.

### Measurement of transcription factors expression by qPCR

Naive, NKG2D− and NKG2D+ memory CD8 T cells were sorted from VV-infected mice (50 days post infection). Total RNA was extracted using TRIzol^®^ reagent according to manufacturer’s instructions (Life Technologies, Saint Aubin, France). Total RNA was digested using turbo DNA-free DNAse (Life technologies, Saint Aubin, France) to avoid genomic contamination. Quality and absence of genomic DNA contamination were assessed with a Bioanalyzer (Agilent, Massy, France). We used High capacity RNA-to-cDNA kit (Life technologies, Saint Aubin, France) to generate cDNA for PCR amplification. PCR was carried out with a SybrGreen-based kit (FastStart Universal SYBR Green Master, Roche, Basel, Switzerland) on a StepOne plus instrument (Applied biosystems, Calrlsbad, USA). Primers were designed using the Roche website (Universal Probe Library Assay Design Center).

### Transwell assay

1 × 10^6^ CD8 T cells purified from VV-infected mice (50 days post infection) were added to the upper chamber of polycarbonate transwell inserts (Corning, 5 μm pore size). The lower chamber was filled with complete DMEM medium (6% FCS, 10 mM HEPES, 50 μM βmercaptoethanol, 50 μg/mL gentamicin, 2 mM L-glutamin, all from Life Technologies) alone or supplemented with CXCL10 (100 ng, Peprotech). After two hours of incubation at 37°C, 7% CO2, transmigrated cells were collected in the lower chamber, washed and stained for CD8, CD44 and NKG2D. Absolute number of transmigrated cells was determined by flow cytometry by adding a known number of fluorescent beads (Flow-Count Fluorosphere, Beckman Coulter). Results are expressed as migration index, which represent the fold increase in the number of transmigrated cells in response to chemoattractant over the non-specific cell migration (medium alone).

### CD3 antibody-directed Cytotoxicity assay

P815 target cells were labelled with CTV (Thermofisher) and incubated with anti-CD3 antibody (2C11, 10 μg/mL) for 30min at 37°C. Target cells were cultured for 12 hours with naive, NKG2D− or NKG2D+ memory CD8 T cells sorted from VV-infected C57BL/6 mice (80 days post-infection). P815 viability was assessed using Live/dead fixable dye (Life technologies) followed by fixation with 1% PFA followed by flow cytometry analysis. The percentage of specific lysis is the percentage of P815 cell death in samples containing CD8 T cells after substracting the percentage of spontaneous P815 cell death.

## Acknowledgments

We thank Dr Wencker for critical reading of the manuscript. We also acknowledge the contribution of SFR BioSciences (UMS3444-CNRS/US8-INSERM ENSL, UCBL) facilities: T. Andrieu and S. Dussurgey (AniRA-Cytométrie) and JL. Thoumas, C. Angleraux and JF. Henry (AniRA-PBES). This work was supported by INSERM, CNRS, Université de Lyon, ENS-Lyon, région Rhône-Alpes and ANR (grant 12-RPIB-0011). MG was a région Rhône-Alpes PhD fellow.

